# Chitin oligomers directly promote lymphoid innate and adaptive immune cell activation

**DOI:** 10.1101/2022.04.06.487356

**Authors:** Yamel Cardona Gloria, Katharina Fuchs, Tzu-Hsuan Chang, Pujan Engels, Elisa Rusch, Cécile Gouttefangeas, Alexander N.R. Weber

## Abstract

**Background:** Chitin is a highly abundant N-acetyl-glucosamine (GlcNAc) polysaccharide which has been linked to immune responses in the context of fungal infections and allergic asthma, especially T helper 2 (Th2) immune responses. Unfortunately, due to the frequent use of crude chitin preparations of unknown purity and degree of polymerization there is still great uncertainty how chitin activates different parts of the human immune system.

**Objective:** We recently identified chitin oligomers of six GlcNAc units as the smallest immunologically active chitin motif and the innate immune receptor TLR2 as primary chitin sensor on human and murine myeloid cells, but the response of lymphoid cells to oligomeric chitin has not been investigated and was addressed here.

**Methods:** Flow cytometric analysis of primary human immune cells upon stimulation with oligomeric chitin. For detailed Methods, please see the Methods section in this article’s Online Repository at www.jacionline.org.

**Result:** Here we report that chitin oligomers also activate immune responses by both primary lymphoid innate and adaptive immune cells: Notably, chitin oligomers activated Natural Killer (NK) cells and B lymphocytes. Moreover, the maturation of dendritic cells enabled CD8 T cell recall responses.

**Conclusions:** Our results suggest that chitin oligomers not only trigger immediate innate responses in limited range of myeloid cells but exert critical activities across the entire human immune system.

**Key messages:** Chitin oligomers are microbe-associated molecular patterns involved in TLR2-dependent fungal recognition by myeloid immune cells.

Here we show that lymphoid cells, most notably Natural Killer (NK) cells and B cells, are activated by oligomeric chitin.

Oligomeric chitin also promotes antigen-presenting cell (APC) maturation and CD8 T cell recall responses.

**Capsule summary:** Oligomeric chitin is a novel microbe-associated molecular patterns involved in TLR2-dependent fungal recognition by myeloid innate immune cells. Here we show that lymphoid innate and adaptive immune responses are also promoted by oligomeric chitin.

## Introduction

Chitin, a β-(1-4)-N-acetyl D-glucosamine (NAG, also abbreviated as GlcNAc) homo-polymer ^1^, is a highly abundant polysaccharide in nature. It is found in the cell walls of fungi, the microfilarial sheaths of parasitic nematodes, the exoskeleton of crustaceans and insects, and as a component of house dust mites - organisms to which vertebrates are exposed through e.g., food ingestion, inhalation or infection ^2, 3^. Its absence in humans makes chitin an ideal target for recognition by the host immune system. In fact, this biopolymer is known to elicit strong immunogenic activity with particular relevance for airway inflammation during asthma ^4, 5^ and fungal infections; pathological conditions that cause socio-economic burden on a pandemic scale ^6, 7^. As a highly conserved microbe-associated molecular pattern (MAMP) ^2^, chitin has been shown to attract and activate innate immune cells and induce cytokine and chemokine production upon recognition by pattern recognition receptors (PRRs) in various cell types, including lung and intestinal epithelial cells and macrophages ^3, 8^. Surprisingly, the immune receptor mediating the observed pleiotropic immunological effects of chitin has long been unclear, due to the repeated use of crude chitin with unknown purity and molecular composition in previous studies ^9^.

In recent work we used purified and defined chitin of known degree of polymerization, i.e. oligomer length, rather than crude chitin, and identified six-subunit-long chitin oligomers as the smallest immunologically active motif. Additionally, we identified the innate immune receptor Toll-like receptor (TLR2) as a primary fungal chitin sensor on human and murine myeloid cells {Fuchs, 2018 #9694} and the host chitinase CHIT1 as an enzyme involved in oligomer generation ^10^. Illustrating the validity of using pure chitin oligomers, He and colleagues have also shown that chitin oligomers can be recognized by the Lysin motif (LysM)-domain containing receptor LYSMD3 on the surface of human airway epithelial cells and elicit production of inflammatory cytokines ^11^. However, so far the effects of oligomeric chitin on lymphoid innate (NK cells) and adaptive (B and T lymphocytes) immune cells have not been addressed.

Here we show that chitin oligomers activate NK cells and human primary B cells, induce maturation of antigen presenting cells such as primary human monocyte-derived dendritic cells (MoDCs) and provide a strong co-activatory signal to antigen-specific CD8^+^ T cells. By shedding light into the effects of chitin oligomers on innate and adaptive immune cells, our work opens new avenues for the development of prophylactic and therapeutic strategies against chitin-mediated inflammatory conditions or the use of chitin oligomers as adjuvants.

## Results and discussion

In order to complement earlier work on myeloid cells, we here tested the effects of chitin oligomers consisting of 10-15 GlcNAc units (termed C10-15, MW range 2,000-3,000 Da, see Methods) on lymphoid innate (NK cells) and adaptive (B and T lymphocytes) primary human immune cells, compared to the well-known TLR2 ligand, Pam_3_CSK_4_ (here termed Pam3) ^12^, and further control stimuli.

Firstly, we assessed CD3^−^ CD56^+^ NK cells in primary PBMC cultures (see Methods) and found that C10-15 led to a robust upregulation of the activation marker CD69 in gated NK cells (Fig. 1A, B). Pam3 was less effective at CD69 upregulation in these cells. This suggests that not only myeloid innate cells can respond to the oligomeric MAMP chitin but also lymphoid innate cells such as NK cells, which are known to express TLR2 ^13^. This warrants further investigation into the effects of chitin oligomers on NK-mediated cytokine production and their capacity as cytotoxic effector cells.

**Figure 1:**
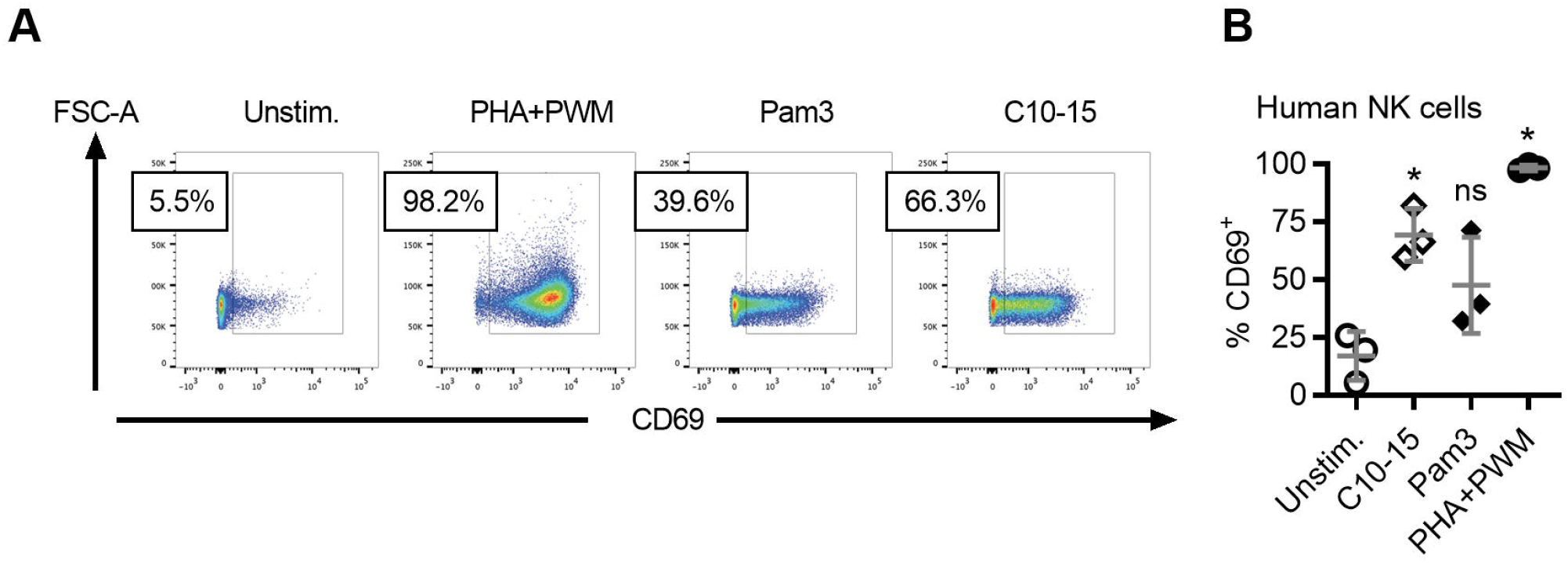
Stimulation of human CD3^−^ CD56^+^ NK cells with chitin oligomers leads to upregulation of the activation marker CD69. (A-B) Fresh PBMCs were cultured in the presence of PHA-L+PWM, Pam3, C10-15 (10 μg/ml) or left unstimulated. After 40 h, the activation status of different populations was assessed by measuring CD69 expression. The following gating strategy was applied: time gate, single cells (FSC-H/FSC-A), living cells (Zombie-Aqua /FSC-A), lymphocytes (FSC-A/SSC-A); NK cells were defined as CD14^−^CD3^−^CD19^−^ CD56^+^ cells. (A) Histograms of a representative donor and (B) summary of CD69 upregulation in gated NK cells within PBMC cultures (n=3 donors). In A and B one representative of three biological replicates are shown. B represents combined data (mean+SD) from ‘n’ biological replicates (each dot represents one donor). * p<0.05 according to one-way ANOVA with Sidak’s correction for multiple testing compared to unstimulated (B).

Next, CD19^+^ B cells within PBMC were analyzed and found to be moderately but significantly activated (CD69 upregulation) by C10-15 compared to Pam3 (Fig. 2A, B). Nevertheless and in line with earlier reports that TLR2 is functional in primary human B cells ^14^, purified CD19^+^ human primary B cells directly responded to C10-15 with IL-6 secretion, albeit to a lower level than to the strong B cell stimulus CpG (Fig. 2C).

**Figure 2:**
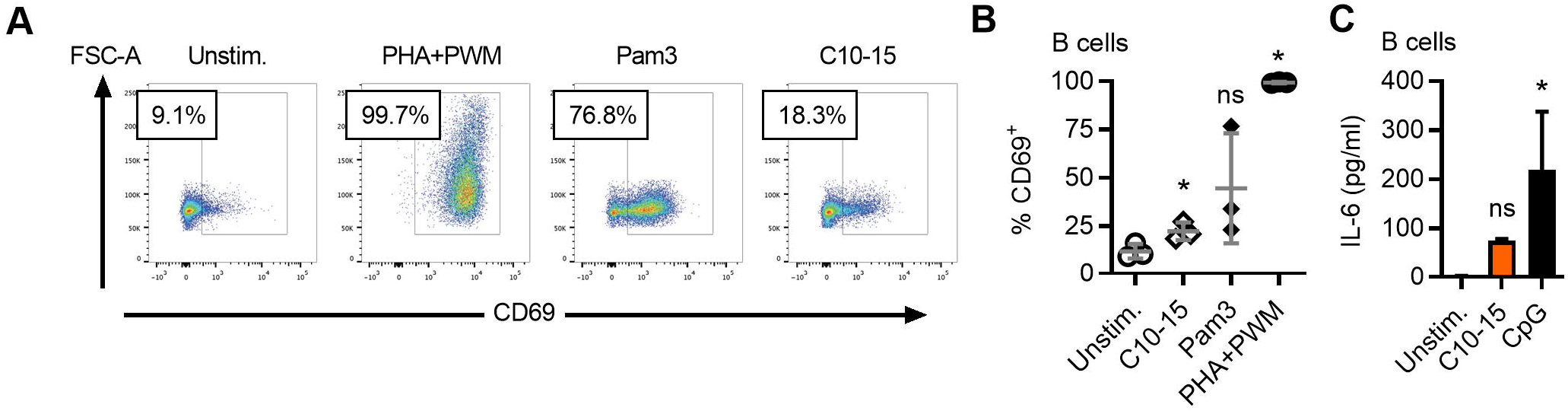
Moderate activation of CD19^+^ human B cells by chitin oligomers. (A) Histograms of a representative donor and (B) summary of CD69 upregulation in gated B cells (CD14^−^ CD3^−^CD19^+^) within PBMC cultures (n=3 donors). (C) IL-6 release from primary purified CD19^+^ human B cells (n=3 donors). In A one representative of three biological replicates is shown. B and C represent combined data (mean+SD) from ‘n’ biological replicates (each dot represents one donor). * p<0.05 according to one-way ANOVA with Sidak’s correction for multiple testing (B) and Friedman test with Dunn’s correction for multiple testing (C), compared to unstimulated.

Conversely, T cells were not directly activated by C10-15 or Pam3, as there was no CD69 upregulation (Fig. 3A) by either TLR2 stimulus. However, in antigen recall experiments using two model HLA class I peptides (EBV-derived BMLF1 and LMP2 sequences), we found that C10-15 could provide a strong and specific co-activatory signal to antigen-specific CD8^+^ T cells. The frequency of BMLF1- and LMP2-specific CD8^+^ T cells was increased of approx. 3-fold (4.5% vs 1.5% and 30.2% vs 10.0% for BMLF1 and LMP2, respectively) in the presence of a very low dose of IL-2 and C10-15 (Fig. 3B, C). In all four analyses (two different peptides with and without IL-2 supplementation) performed with cells from two donors C10-15 showed this effect. This was confirmed in another donor; however, here the condition using LMP2 peptide +0.2 ng/ml IL-2 did not show an effect for C10-15 (not shown), for unknown reasons, potentially the occurrence of a hypofunctional *TLR2* allele {Fuchs, 2018 #9694}. It is noteworthy that C10-15 chitin oligomers were more potent than the lipopeptide Pam3 on which novel adjuvants recently applied to a human SARS-CoV2 T cell vaccine are based {Rammensee, 2019 #7688}{Heitmann, 2022 #10996}. Overall, chitin oligomers can indirectly but potently promote antigen-dependent CD8 T cell responses.

**Figure 3:**
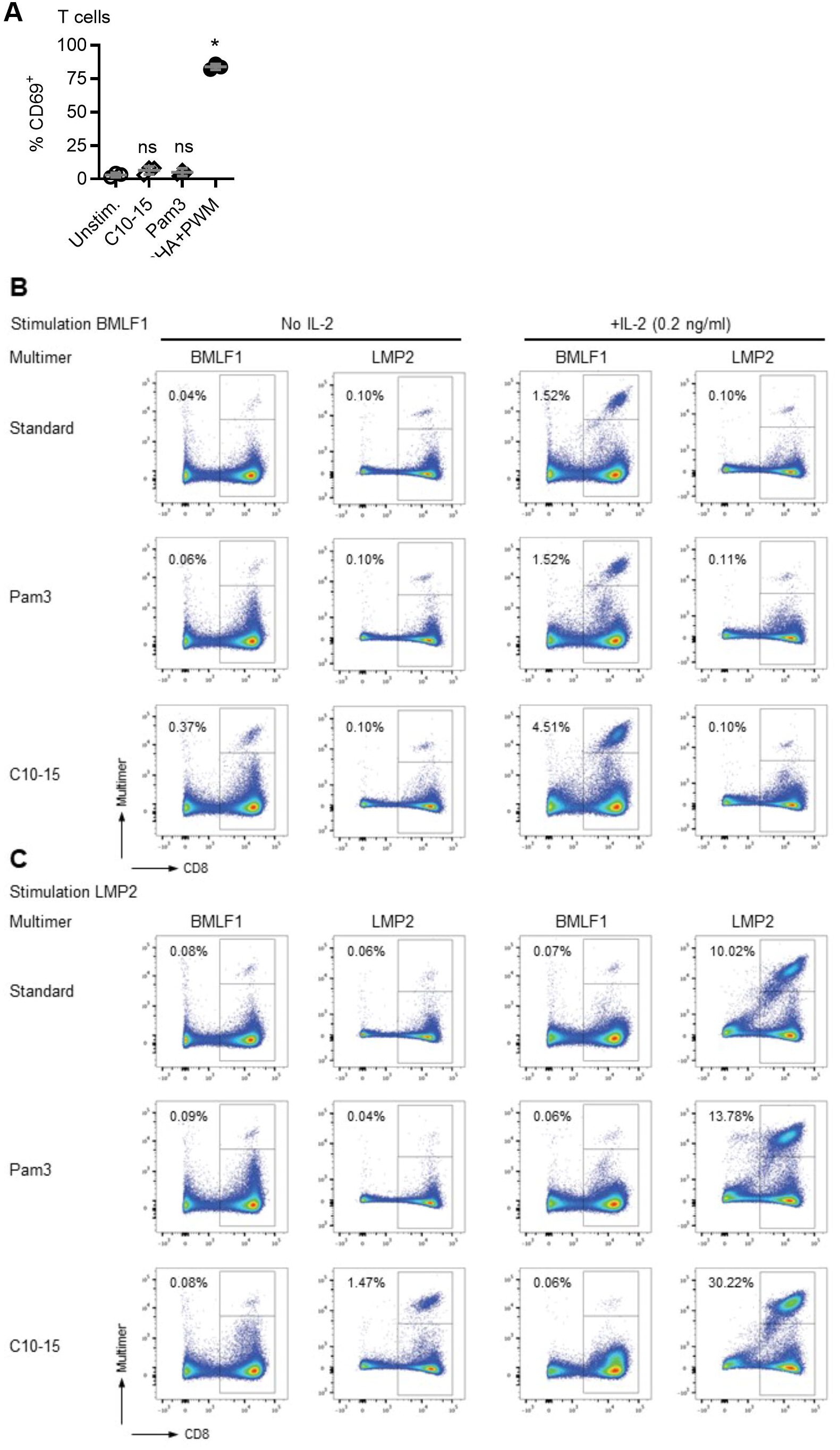
Effect of chitin oligomers on T lymphocytes. (A) CD8^+^ multimer^+^ T cells expanded from human PBMC (n=2) in the presence of antigenic peptides, with or without addition of a low dose of recombinant hIL-2: PBMCs from one representative donor were cultured for 12 days in the presence of BMLF1 (A, upper panels) or LMP2 (B, lower panels) antigenic peptides and either no (left column) or 0.2 ng/ml (right column) r-hIL-2; either standard medium was used or Pam3 or C10-15 were added to 10 μg/ml. Cells were harvested and stained with HLA-A*02 multimers, anti-CD4-FITC, anti-CD8-PE-Cy7 antibodies and Zombie-Aqua. Columns 1 and 3 show the staining with the BMLF-1 tetramer, columns 2 and 4 show staining with the LMP2 tetramer. The following gating strategy was applied: time gate, single cells (FSC-H/FSC-A), living cells (Zombie Aqua /FSC-A), lymphocytes (FSC-A/SSC-A), CD4^−^ (CD4/CD8); pseudocolor plots show CD8 (x-axis) vs multimer staining (y-axis). Percentages of multimer^+^ CD8^+^ cells are given for each single staining performed in 1 replicate, one out of two tested donors is shown. B represents combined data (mean+SD) from ‘n’ biological replicates (each dot represents one donor). (C) Summary of CD69 upregulation in gated T cells (CD14^−^CD3^+^CD19^-^) within PBMC cultures (n=3 donors). * p<0.05 according to one-way ANOVA with Sidak’s correction for multiple testing compared to unstimulated (C).

Based on the fact that T cells did not respond directly in short-term cultures (*cf*. Fig. 3A), we hypothesized that chitin oligomers acted on antigen presenting cells, e.g. dendritic cells, contained in the PBMC. This hypothesis was congruent with the observation that C10-15 led to potent maturation of primary human monocyte-derived dendritic cells (MoDCs), a type of antigen presenting cells, as indicated by HLA-DR, CD83 and CD86 upregulation (Fig. 4A-C). In general, C10-15 is able to strongly activate antigen presenting cells, which support/evoke efficient and specific CD8^+^ T cell responses and more than Pam3. Collectively, chitin oligomers thus trigger immune responses in both myeloid human immune cells {Fuchs, 2018 #9694} and lymphoid innate cells (NK cells) and adaptive immune cells (B cell and T cells)

**Figure 4:**
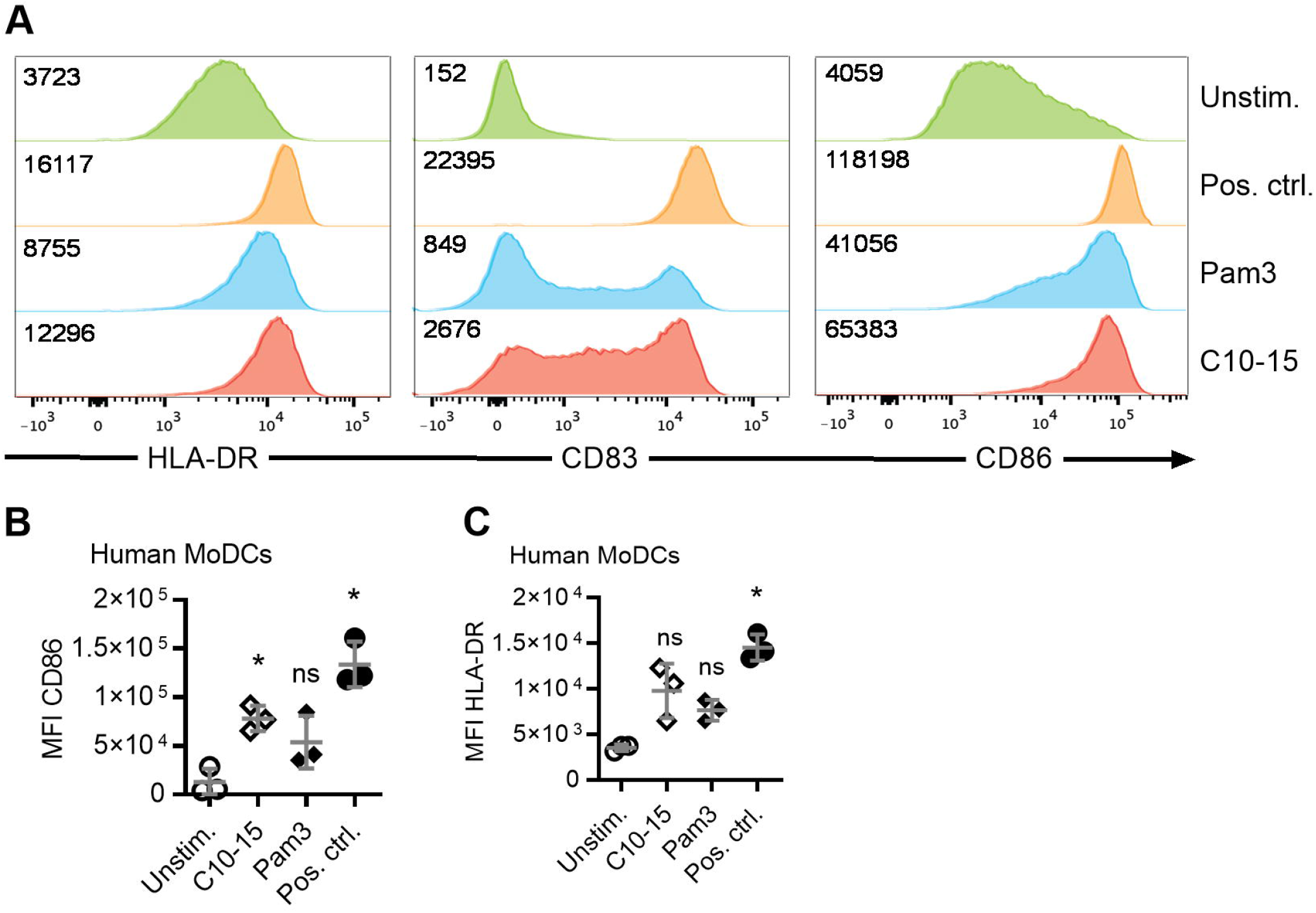
Chitin oligomers lead to potent maturation of primary human monocyte-derived dendritic cells (MoDCs). Upregulation of maturation markers in human MoDCs for one representative (A) out of n=3 donors (B and C), with median fluorescence intensities indicated in (A). B (CD86) and C (HLA-DR) represent combined data (mean+SD) from ‘n’ biological replicates (each dot represents one donor). * p<0.05 according to one-way ANOVA with Dunnett’s correction for multiple testing compared to unstimulated (B), and Friedman test with Dunn’s correction for multiple testing (C), compared to unstimulated.

Collectively, chitin oligomers appear to have broad adjuvant-like properties, i.e. activation of multiple different human innate immune cell types and stimulation of adaptive immune responses. Compared to earlier results, chitin oligomers thus not only possess a strong capacity to fully signal danger to epithelial ^11^ and myeloid innate cell types (e.g. macrophages, PMNs)^10, 15^ but also to lymphoid innate immune cells such as NK cells. They can also directly act on B lymphocytes in an antigen-independent manner. B lymphocytes with an antigen specificity for chitin have been described in more severe cases of inflammatory bowel diseases ^16^. It could thus be conceivable that in certain individuals and for certain B cell clones, chitin oligomers may not only activate TLR2 but also engage BCR signaling for potent B cell activation. By activating antigen-presenting cells such as MoDCs, chitin oligomers are also able to indirectly promote T cell mediated adaptive immunity and display adjuvant like properties. It could be envisaged that in house dust mite allergies, chitin oligomers, once released from *Dermatophagoides* house dust mites by the host chitinase, CHIT1 ^10^, thus perform a “self-adjuvanting” role for associated T cell antigens ^17^. From a translational perspective, our study raises the possibilities that blocking chitin-TLR2 interactions might prevent immune activation more broadly than previously thought or that chitin oligomers could be a useful addition to the short list of adjuvants suitable for use in humans ^18^.

## Abbreviations

(CLR): C-type lectin receptor
(ELISA): enzyme-linked immunosorbent assay
(IL): interleukin
(LPS): lipopolysaccharide
(MFI): median fluorescence intensity
(MAMP): microbe-associated molecular pattern
(MyD88): myeloid differentiation primary response 88
(NAG, GlcNAc): N-acetyl-glucosamine
(NK cell): Natural Killer cell
(NF-κB): nuclear factor ‘kappa-light-chain-enhancer’ of activated B-cells
(PRRs): pattern recognition receptors
(PBMCs): peripheral blood mononuclear cells
(PMN): polymorphonuclear neutrophil
(TLR): Toll-like receptor
(TNF): Tumor necrosis factor

## Acknowledgements

We thank Sabine Dickhöfer for excellent technical assistance and Libera Lo Presti for editorial support.

## Author contributions

YCG, KF, THC, PE, ER performed experiments; YCG, KF, ER, PE, CG, AW analyzed data; AW wrote the manuscript which all other authors commented on; AW conceived and coordinated the study.

## Methods Summary

### Reagents and antibodies

All chemicals were from Sigma unless otherwise stated. C10-15 chitosan oligomers (a mixture of fragments in an MW range of 2000-3000) were from Carbosynth. Chitin C10-15 was generated from chitosan oligomers using sodium bicarbonate and acetic anhydride acetylation ^19^. Purities of >95%, acetylation of >90% were achieved as described in Supplemental Information. Pam3 was from Invivogen, CpG 2006 from TIB Molbiol and, LMP2 and BMLF1 peptides synthesized in house. Polymyxin B was from ThermoFisher.

### Isolation of PBMC from human peripheral blood

Peripheral blood mononuclear cells (PBMCs) were isolated from heparinized whole blood or buffy coats from healthy human volunteers by standard density gradient separation (Biocoll, Biochrom), washed twice in PBS and counted with trypan blue. Cells were used either immediately or frozen in FCS containing 10% DMSO and transferred to liquid nitrogen until use.

### Isolation and analysis of primary human B cells

Following PBMC preparation, B cells were isolated using anti-CD19 microbeads (Miltenyi Biotec) reaching >90% purity (as assessed by anti-CD19-PE positivity). Upon isolation primary B cells were seeded in RPMI supplemented with 20% heat inactivated fetal calf serum (FCS), rested for at least 4 h and treated for 18 h with C10-15 or CpG 2006 at 5 µM. Cytokines in supernatants were measured by ELISA using a human IL-6 ELISA kit (Biolegend).

### Isolation and analysis of human dendritic cells

Monocyte-derived dendritic cells were obtained by plate adherence of PBMCs following a 3-day procedure of differentiation with GM-CSF (1000 U/ml) and IL-4 (25 U/ml, both PeproTech). After 48 h, cells were either left untreated or matured with Pam3 or C10-15 (each 10 µg/ml). Twenty four hours later, cells were harvested, Fc blocked (BD Biosciences), stained with anti-CD14 Alexa-Fluor700 (eBioscience, clone 61D3), -CD83-APC (Biolegend, clone HB15e), -HLA-DR-PerCP (BD, clone L243), CD86-BV605 (Biolegend, clone IT2.2) and Zombie Aqua (Biolegend), measured on a LSR Fortessa (BD) and analyzed using FlowJo PC version 10. Further details see Supplemental Information.

### Human B and NK cell activation assay

Freshly isolated PBMCs were cultured in supplemented IMDM medium either alone or with C10-15, Pam3 (each 10 µg/ml), or a mix of phytohemagglutinin-L and Pokeweed mitogen (2 and 1 µg/ml, respectively). After 40 h, cells were harvested, stained with anti-CD3-BV711 (Biolegend, clone OKT3), -CD19-BV785 (Biolegend, clone HIB19), -CD14-Alexa700, -CD56-BV421 (Biolegend, clone HCD56), - HLA-DR-PerCP, -CD69-APC-Cy7 (BD, clone FN50) mAbs and Zombie Aqua (Biolegend). Acquisition was performed as described above. Further details see Supplemental Information.

### Expansion and analysis of human antigen-specific T cells

PBMC from an HLA A*02 positive donor (with *ex vivo* detectable EBV LMP2- and BMLF1-specific CD8^+^ T cells) were incubated with C10-15, Pam3 (each 10 µg/ml) or medium only in combination with peptides (1 µg/ml of EBV LMP2 426-434 CLGGLLTMV or EBV BMLF1 259-267 GLCTLVAML, both known HLA-A*02 T cell epitopes). Cultures were either not supplemented or supplemented with 0.2 ng/ml of recombinant human IL-2 (rIL-2 R&D Systems) at days 3, 5, 7 and 9, and split if needed. At day 12, cells were harvested and stained with relevant HLA-multimers, anti-CD4-FITC (clone HP2/6, in-house production) and anti-CD8-PE-Cy7 (clone SFCI21-Thy2D3, Beckman Coulter) antibodies and Zombie-Aqua as described in Supplemental Information.

### Statistical analysis

Experimental data was analyzed using Excel 2010 (Microsoft) and/or GraphPad Prism 8, flow cytometry data using FlowJo software version 10. Normal distribution of data was assessed using Shapiro-Wilk test and determined the choice of subsequent parametric or non-parametric tests as indicated in the figure legends. p-values were determined (α=0.05, β=0.8) as indicated and multiple comparisons were corrected for. p-values < 0.05 were generally considered statistically significant and are denoted by * throughout even if calculated p-values were lower than p=0.05.

## References

1. Morgulis S. The Chemical Constitution of Chitin. Science 1916; 44:866–7.

2. Lee CG, Da Silva CA, Lee JY, Hartl D, Elias JA. Chitin regulation of immune responses: an old molecule with new roles. Current opinion in immunology 2008; 20:684–9.

3. Mack I, Hector A, Ballbach M, Kohlhaufl J, Fuchs KJ, Weber A, et al. The role of chitin, chitinases, and chitinase-like proteins in pediatric lung diseases. Mol Cell Pediatr 2015; 2:3.

4. Choi JP, Lee SM, Choi HI, Kim MH, Jeon SG, Jang MH, et al. House Dust Mite-Derived Chitin Enhances Th2 Cell Response to Inhaled Allergens, Mainly via a TNF-alpha-Dependent Pathway. Allergy Asthma Immunol Res 2016; 8:362–74.

5. Van Dyken SJ, Liang HE, Naikawadi RP, Woodruff PG, Wolters PJ, Erle DJ, et al. Spontaneous Chitin Accumulation in Airways and Age-Related Fibrotic Lung Disease. Cell 2017; 169:497–509 e13.

6. To T, Stanojevic S, Moores G, Gershon AS, Bateman ED, Cruz AA, et al. Global asthma prevalence in adults: findings from the cross-sectional world health survey. BMC Public Health 2012; 12:204.

7. Brown GD, Denning DW, Gow NA, Levitz SM, Netea MG, White TC. Hidden killers: human fungal infections. Sci Transl Med 2012; 4:165rv13.

8. Lee CG, Da Silva CA, Lee JY, Hartl D, Elias JA. Chitin regulation of immune responses: an old molecule with new roles. Curr Opin Immunol 2008; 20:684–9.

9. Bueter CL, Specht CA, Levitz SM. Innate sensing of chitin and chitosan. PLoS Pathog 2013; 9:e1003080.

10. Chang T-H, Gloria YC, Hellmann MJ, Greve CL, Roy DL, Roger T, et al. Transkingdom mechanism of MAMP generation by chitotriosidase (CHIT1) feeds oligomeric chitin from fungal pathogens and allergens into TLR2-mediated innate immune sensing. bioRxiv 2022:2022.02.17.479713.

11. He X, Howard BA, Liu Y, Neumann AK, Li L, Menon N, et al. LYSMD3: A mammalian pattern recognition receptor for chitin. Cell Rep 2021; 36:109392.

12. Grabiec A, Meng G, Fichte S, Bessler W, Wagner H, Kirschning CJ. Human but not murine toll-like receptor 2 discriminates between tri-palmitoylated and tri-lauroylated peptides. J Biol Chem 2004; 279:48004–12.

13. Lauzon NM, Mian F, MacKenzie R, Ashkar AA. The direct effects of Toll-like receptor ligands on human NK cell cytokine production and cytotoxicity. Cell Immunol 2006; 241:102–12.

14. Bekeredjian-Ding I, Jego G. Toll-like receptors--sentries in the B-cell response. Immunology 2009; 128:311–23.

15. Fuchs K, Cardona Gloria Y, Wolz OO, Herster F, Sharma L, Dillen CA, et al. The fungal ligand chitin directly binds TLR2 and triggers inflammation dependent on oligomer size. EMBO Rep 2018; 19.

16. Rieder F, Schleder S, Wolf A, Dirmeier A, Strauch U, Obermeier F, et al. Association of the novel serologic anti-glycan antibodies anti-laminarin and anti-chitin with complicated Crohn’s disease behavior. Inflamm Bowel Dis 2010; 16:263–74.

17. Oseroff C, Christensen LH, Westernberg L, Pham J, Lane J, Paul S, et al. Immunoproteomic analysis of house dust mite antigens reveals distinct classes of dominant T cell antigens according to function and serological reactivity. Clin Exp Allergy 2017; 47:577–92.

18. Khong H, Overwijk WW. Adjuvants for peptide-based cancer vaccines. J Immunother Cancer 2016; 4:56.

19. Bueter CL, Lee CK, Rathinam VA, Healy GJ, Taron CH, Specht CA, et al. Chitosan but not chitin activates the inflammasome by a mechanism dependent upon phagocytosis. J Biol Chem 2011; 286:35447–55.

